# Spatio-temporal mapping of immune cell dynamics during human sequential lymph node metastasis

**DOI:** 10.64898/2026.03.17.712495

**Authors:** Qiuchen Zhao, Yongjin Lu, Zhiqiang Shi, Hongrun Zhang, Li Cheuk Shuen, Rongrong Zhao, Yuanchao Ling, Yongsheng Gao, Zhaopeng Zhang, Xiao Sun, Yuping Qian, Xiaowen Wang, Chunjian Wang, Binbin Cong, Xiang Ni, Yanfang Liu, Miaoqing Zhao, Yongsheng Wang, Bidesh Mahata, Pengfei Qiu

## Abstract

Regional lymph node (LN) metastasis critically influences distant metastatic progression, anti-tumour immunity, and patient prognosis. While tumour-induced immune modulation in tumour-draining LNs (TDLNs) has been extensively studied using murine models, the systematic reconstruction of the immune system from primary tumours through TDLNs and subsequent lymph nodes in human cancer progression remains understudied. Here, we utilised integrated multi-omics approaches, including imaging mass cytometry, single-cell RNA sequencing, Visium and Xenium spatial transcriptomics, and multi-colour immunofluorescence to systematically characterise immune cell dynamics across 147 paired primary tumours, sentinel TDLNs (S-TDLNs), and secondary axillary LNs (ALNs) obtained from 50 treatment-naïve triple-negative breast cancer patients with different progression statuses. Our comprehensive profiling revealed critical immune alterations, such as decreased type-2 conventional dendritic cells (cDC2), naïve T cells, and B cells, along with an increase in immunosuppressive macrophages. Developing a novel single-cell transformer model, we identified substantial alterations in various immune cell populations, notably MARCO^+^ macrophages, which strongly correlated with breast cancer patient survival outcomes. Spatial analysis combined with our newly integrated cell-cell interaction platform revealed diminished immune cell communication and impaired priming interactions among dendritic cells, B cells, and T cells within metastatic lymph nodes and primary tumour sites. In an independent neoadjuvant immunotherapy cohort of 36 TNBC patients with 52 lymph node samples, we found preservation of CD1c⁺ cDC2 in lymph nodes predicted pathological complete response and longer event-free survival, highlighting cDC2 as a potential biomarker and therapeutic target. Collectively, this systematic mapping of immune landscape alterations during human sequential LN metastasis provides essential insights for understanding cancer metastasis mechanisms and paves the way for innovative immunotherapeutic strategies.

## INTRODUCTION

Regional lymph node (LN) metastasis is an important event in cancer progression, playing roles in distant metastatic process, anti-tumour immunity, and clinical outcomes ^1,2^. Tumour-draining lymph nodes (TDLNs), as the primary sites of tumour-immune interactions, undergo profound immune modulation that influences both local and systemic anti-tumour responses ^3–5^. Some studies in murine models have showed the cellular and molecular mechanisms by which tumours educate TDLNs to foster immune tolerance, modulate T cell subsets, and alter the stromal microenvironment ^6–9^. However, the findings are derived mainly from animal studies or unpaired clinical samples, with limited resolution at the single-cell or spatial resolution in human tissues ^10^. Systematic, high-dimensional reconstruction of immune dynamics from primary tumours through sequential lymph node stations in human cancer remains rare, largely due to the challenges in obtaining well-annotated, multi-omic paired samples across the metastatic trajectory.

Breast cancer provides an ideal clinical model to investigate the stepwise immune and metastatic changes in regional lymph nodes, because of its globally practice of sentinel lymph node biopsy (SLNB), a milestone in minimally invasive oncologic surgery ^11^. Over the past decades, axillary management in breast cancer has shifted towards de-escalation, with increasing numbers of patients undergoing only SLNB or even being spared lymph node dissection entirely, thereby preserving TDLNs for potential study ^12–17^. Despite these advances, current clinical settings primarily focus on the presence or absence of tumour cell metastasis, treating lymph nodes as passive tubes rather than active hubs of immune regulation. The sentinel lymph node, as the first regional node exposed to tumour-derived factors, is uniquely positioned to reflect early immune reprogramming and may critically influence both local and systemic anti-tumour immunity. This is particularly relevant in highly immunogenic but therapeutically refractory subtypes such as triple-negative breast cancer (TNBC), where failure to consider dynamic immune modulation within lymph nodes may lead to missed opportunities for precisely timed immunotherapeutic intervention.

TNBC is characterised by high immunogenicity, substantial intra-tumoural heterogeneity, and poor clinical outcomes ^18,19^. Although TNBC tumours exhibit elevated infiltration of cytotoxic T cells and other immune effectors, they are frequently accompanied by profound immunosuppressive mechanisms, including expansion of regulatory T cells, myeloid-derived suppressor cells, and exhaustion of effector lymphocytes within the tumour microenvironment ^20–24^. These immune escape pathways are well documented at the primary tumour site, where immune checkpoint blockade therapies have shown only limited benefit despite the immunogenic nature of TNBC ^19^. However, the high-resolution immune landscape of tumour-draining and regional lymph nodes in TNBC remains less explored, with most studies to date focused on tumour tissues rather than the nodal microenvironment ^25–27^. Recent evidence suggests that the immunological profile of lymph nodes can both shape systemic anti-tumour responses and serve as a reservoir for immune evasion, thereby directly impacting patient prognosis and therapeutic efficacy ^28^. As a result, the extent to which immunosuppressive microenvironments are established, maintained, or reshaped within sentinel TDLNs (S-TDLNs), and secondary axillary LNs (ALNs) during metastatic progression of TNBC is critical to explore.

TNBC represents only 10–15% of breast cancers and is frequently treated with neoadjuvant therapy, limiting access to treatment-naive samples and metastatic LNs ^29^. The adoption of sentinel lymph node biopsy has further reduced the availability of non-sentinel lymph nodes ^30^. As a result, human studies capturing the full immune trajectory from primary tumours through sentinel and downstream nodes remain scarce.

Here, we assembled a unique cohort of spatially and transcriptionally profiled, treatment-naive TNBC specimens across different stages of nodal metastasis, enabling a comprehensive reconstruction of the immune landscape across tumour, S-TDLNs, and secondary lymph nodes. Our study aims to explore the stepwise immune reprogramming events and disruption of lymphoid architecture during LN metastasis, with particular focus on identifying key cellular interactions and spatial organisation patterns that drive immune evasion. Through development of single-cell transformer model and integrative cell-cell communication framework, we seek to uncover previously unrecognised immune cell states and communication networks that are disrupted during sequential LN colonisation. Furthermore, by linking our immune landscape to an independent neoadjuvant immunotherapy cohort, we identify CD1c⁺ cDC2 in lymph nodes as a potential predictive biomarker for treatment response and prognosis in TNBC. These findings provide essential mechanistic insights into the immunological basis of lymph node metastasis and inform the development of targeted immunotherapeutic strategies that exploit the preserved immunological functions within tumour-draining lymph nodes to enhance anti-tumour immunity.

## RESULTS

### Early immune remodeling across primary tumours and tumour-draining lymph nodes

We enrolled 50 treatment-naïve TNBC patients to investigate immune modulation during LN metastasis (Table S1, Figure S1A). Of these patients, 20 showed no metastatic involvement in either their sentinel TDLNs or axillary LNs, 15 had metastases in their S-TDLNs but not their ALNs, and the remaining 15 had metastases in both S-TDLNs and ALNs, indicative of more advanced disease progression (Table S1, Figure 1A). We subsequently performed a comprehensive immune profiling using multiple platforms: (1). Imaging mass cytometry (IMC) on 31 FFPE samples (paired primary tumours, S-TDLNs, and ALNs) for immune phenotyping and characterisation of immune cell dynamics, (2). Single-cell RNA sequencing (scRNA-seq) on 29 fresh surgically resected, paired tissues to construct an immune atlas, (3). Visium and Xenium-based spatial transcriptomics on 16 frozen or FFPE samples to examine immune cell distribution and interactions in a spatial context, and (4). Multi-colour immunofluorescence (mIF) on 114 FFPE samples to correlate key immune parameters with clinical outcomes (Figure 1A).

**Figure 1.**
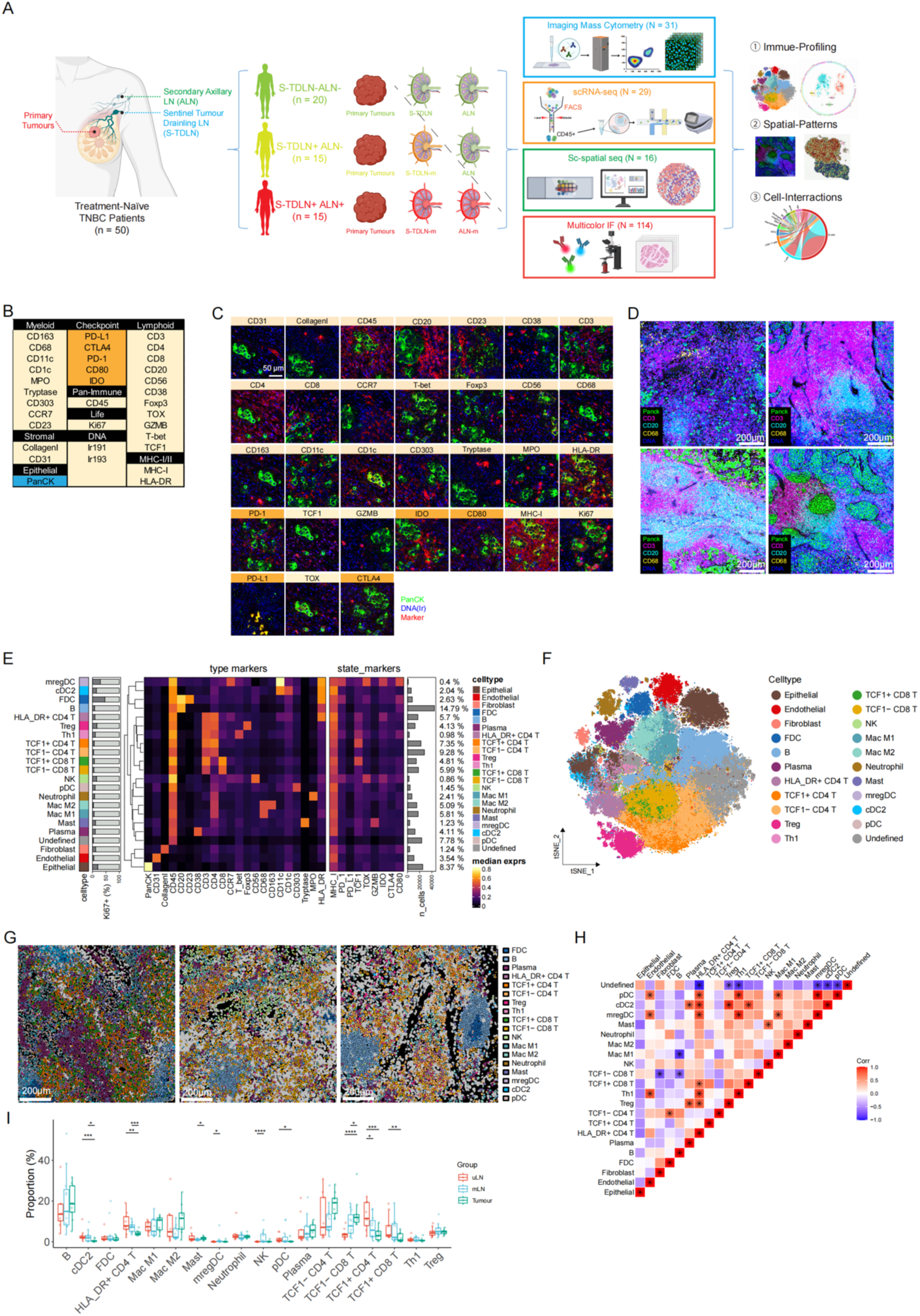
Early immune remodeling across primary tumours and tumour-draining lymph nodes. A, Schematic overview of study design and patient cohorts. Fifty treatment-naïve TNBC patients were stratified into three groups based on nodal metastasis status: no metastatic involvement (STDLN–ALN–, n=20), sentinel-node positive but axillary-node negative (S-TDLN+ALN–, n=15), and both sentinel- and axillary-node positive (S-TDLN+ALN+, n=15). Multi-platform immune profiling, including IMC (n=31), scRNA-seq (n=29), spatial transcriptomics (Visium and Xenium, n=16), and mIF (n=114), was conducted on matched primary tumours, S-TDLNs, and ALNs. B, Antibody panel list targeting 34 protein markers expressed by epithelial cells (blue), TME cells (yellow), or related to immune checkpoint (orange). C, Images of indicated protein markers expression (scale bar 50μm). D, Representative IMC images of primary tumour, sentinel, and axillary lymph nodes stained with antibodies targeting PanCK (tumour epithelial cells), CD3 (T cells), CD20 (B cells), CD68 (macrophages), and DNA. E, Expression heatmap summarising median marker intensities across identified cell types in IMC data, highlighting robust detection of immune, stromal, and epithelial markers. F, t-SNE plot visualising the distinct clustering of immune and non-immune cell populations identified from IMC single-cell data across tissues. G, Spatial distribution of immune cell populations across ALN (left), S-TDLN (middle), and tumour (right) tissues. H, Heatmap illustrating pairwise spatial correlation coefficients between immune cell populations across tumour tissues. I, Quantitative comparison of cell-type proportions among primary tumours, metastatic lymph nodes (mLNs), and uninvolved lymph nodes (uLNs). Data shown as mean ± SEM; statistical significance by one-way ANOVA (*P < 0.05, **P < 0.01, ***P < 0.001, ****P < 0.0001).

Using tissue microarrays (TMAs), we stained tissues with a panel of 34 isotope-labelled antibodies targeting major immune cell phenotypes, functional markers, epithelial, and stromal components (Figure 1B). As expected, we observed robust marker expression (Figure 1C) and distinct immune and cancer cell clusters in tumours, S-TDLNs, and ALNs (Figure 1D). The raw IMC images were then converted into single-cell data for annotation purposes. In total, 21 cell phenotypes were identified based on classical markers, including T cells, B cells, natural killer (NK) cells, macrophages, dendritic cells (DCs), neutrophils, plasma cells, mast cells, epithelial cells, endothelial cells, and fibroblasts (Figure 1E,F, S1A). Among these, some subpopulations were classified by additional markers: FOXP3 for regulatory T cells (Tregs), T-bet for type 1 helper T cells (Th1), TCF1 for stem-like T cells, CD163 for M2 macrophages, CD1C for type-2 conventional DCs (cDC2), CD303 for plasmacytoid DCs (pDCs), and CD80/IDO for mature regulatory DCs (mregDC) (Figure 1E, S1B). We also displayed the spatial distribution of these cell types on slides from ALNs, S-TDLNs, and tumours, identifying clusters of specific immune populations and their interactions with other cell types (Figure 1G).

We further quantified the relative proportions of immune cells in primary tumours, metastatic LNs (mLNs), and uninvolved LNs (uLNs). Notably, cDC2, pDCs, HLA-DR+ CD4+ T cells, and TCF1+ CD4+/CD8+ T cells decreased from uLNs to mLNs and tumours, while M2 macrophages and TCF1– CD4+/CD8+ T cells increased in tumours compared to LNs (Figure 1G). The spatial correlation between cell types were then calculated, and the interactions between cDC2 and Tregs was increased in tumours compared with LNs (Figure 1H, S1C). Together, IMC provides a comprehensive view of spatial distribution, abundance, and neighbourhood patterns of immune populations across disease progression.

### Progressive immune reprogramming and suppression during lymph node metastasis

To construct a detailed single-cell immune atlas and characterise immune cell dynamics during the early stages of LN metastasis, we employed fluorescence-activated cell sorting (FACS) to isolate CD45+ immune cells from 29 fresh, surgically resected primary tumours, S-TDLNs, and ALNs collected from 10 treatment-naïve TNBC patients, followed by single-cell RNA sequencing (scRNA-seq) (Figure 1A, S2A). After rigorous quality control, we obtained single-cell transcriptomic profiles for 246,703 high-quality immune cells, encompassing T cells, B cells, myeloid cells, NK cells, and a cluster of proliferating cells (Figure 2A). Unbiased clustering identified diverse immune cell subtypes within each major immune population, including LEF1+ naïve T (Tn) cells, ANXA1+ central memory CD4+ T (Tcm) cells, GZMK+ effector memory CD8+ T (Tem) cells, FOXP3+ regulatory T (Treg) cells in the T-cell compartment; TCL1A+ naïve B (Bn) cells, CD27+ memory B (Bmem) cells, RGS13+ germinal centre (GC) B cells, and plasma cells in the B-cell lineage; CD1C+ conventional type-2 dendritic cells (cDC2), CCR7+ mature regulatory DCs (mregDCs), LILRA4+ plasmacytoid DCs (pDCs), MARCO+ macrophages, and FCGR3B+ neutrophils among the myeloid population; and FCGR3A+ (CD16+), NCAM1+ (CD56+), KIT+, and CXCL13+ NK cells (Figure 2B, S2B).

**Figure 2.**
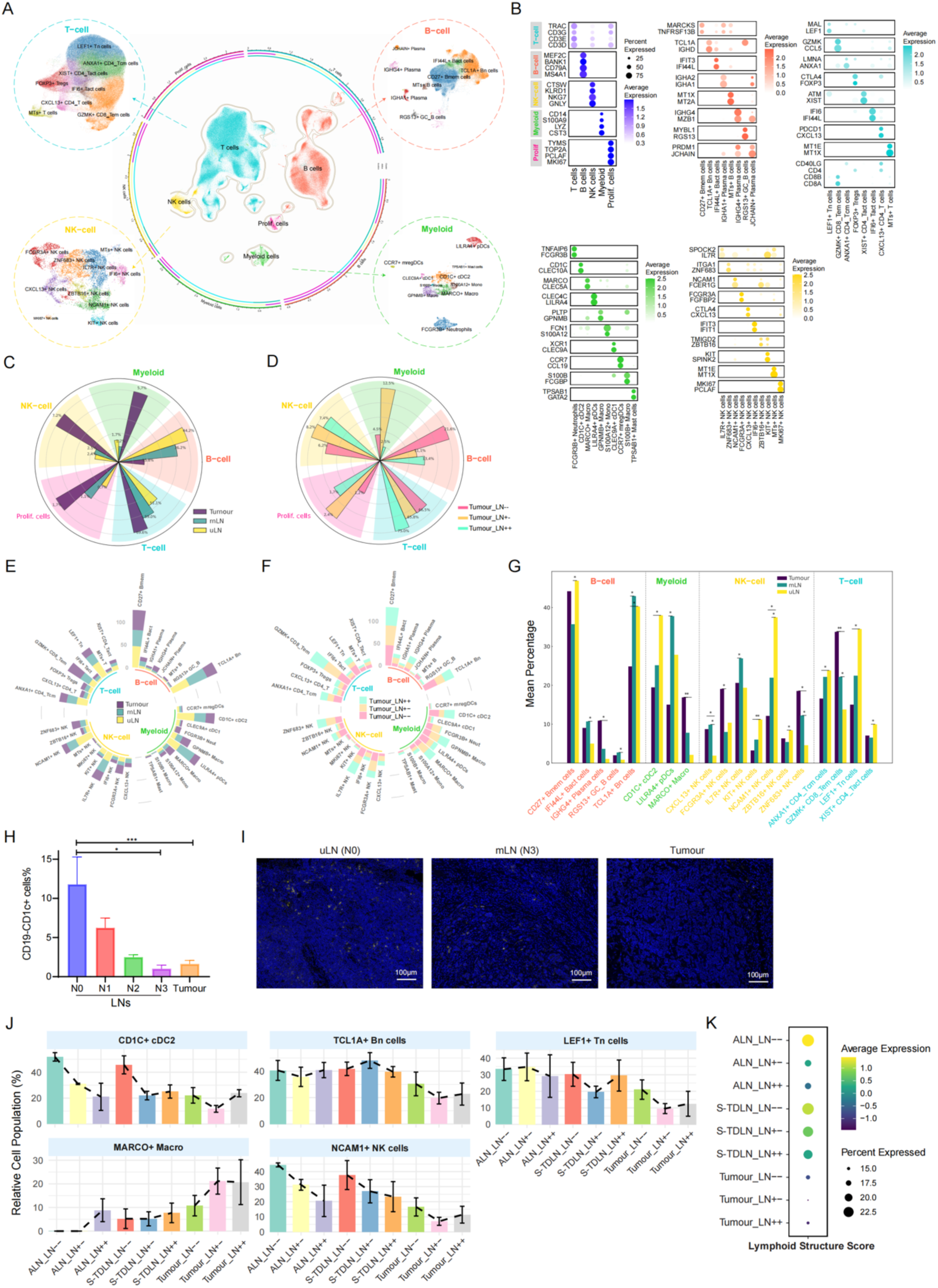
Single-cell immune atlas reveals progressive immune-cell remodelling associated with lymph node metastasis. A, UMAP visualisation of 246,703 high-quality immune cells obtained by scRNA-seq. Major immune cell lineages including T cells, B cells, NK cells, myeloid cells, and proliferating cells are indicated by distinct colours. B, Dot plot depicting canonical marker gene expression defining immune cell populations. Dot size indicates the percentage of cells expressing each gene; colour intensity represents average expression. C, Circular barplot comparing the relative abundance of major immune cell types across tissues (tumours vs. lymph nodes). D, Circular barplot comparing the relative abundance of major immune cell types in tumour tissues across groups (S-TDLN–ALN, S-TDLN+ALN–, and S-TDLN+ALN+). E, Circular barplot showing the relative abundance of 36 immune sub-clusters in each major cell type identified across tumours, mLNs, and uLNs. F, Circular barplot comparing the relative frequencies of immune subpopulations in tumour tissues across nodal metastatic groups (S-TDLN−ALN−, S-TDLN+ALN−, S-TDLN+ALN+). G, Bar plot highlighting significant shifts in immune subpopulations in tumours compared to LNs. H–I, Multi-colour immunofluorescence (mIF) analysis validating the relative frequency of CD19-CD1c+ cDC2 across nodal metastatic stages (N0 to N3) and in tumour tissues (H), with representative images from three tissues illustrated in (I). Scale bar, 100 μm. J, Relative frequencies of selected immune subpopulations across different tissues and metastatic groups. K, Quantification of gene programmes related to lymphoid structure formation across tissues.

Among the major immune cell types, T cells, myeloid cells, and NK cells were relatively more abundant in tumours, whereas B cells showed the opposite trend (Figure 2C). Notably, we also observed a substantial increase in myeloid cells within primary tumours of patients who had S-TDLN metastasis but no ALN involvement (S-TDLN+ALN−), compared with those who lacked metastasis (S-TDLN−ALN−) or had both S-TDLN and ALN metastases (S-TDLN+ALN+), indicating immune modulation during early metastasis (Figure 2D). To clarify the relative proportions of immune subpopulations across different patient groups, we quantified 36 sub-clusters within T cells, B cells, myeloid cells, and NK cells in tumours, mLNs, and uLNs (Figure 2E). We also compared these subpopulations in tumours from the S-TDLN−ALN−, S-TDLN+ALN−, and S-TDLN+ALN+ groups (Figure 2F). In line with our observations of major cell types, several macrophage and neutrophil subclusters were enriched in the S-TDLN+ALN− group (Figure 2F,D). Focusing on the most significant shifts, we noted a marked decrease in TCL1A+ Bn cells, CD1C+ cDC2, NCAM1+ NK cells, and LEF1+ Tn cells in tumours relative to mLNs and uLNs. Conversely, MARCO+ macrophages, ZNF683+ NK cells, and GZMK+ CD8+ Tem cells were increased in tumours (Figure 2G). To further validate these findings and assess their clinical relevance, mIF was employed to examine immune subpopulations across patient samples with known clinical status. Notably, cDC2 frequency progressively declined with advancing nodal involvement in TNBC patients and remained at consistently low levels in tumour tissues (Figure 2H,I, &S2C,D).

Immunocyte populations essential for lymphoid architecture and tertiary lymphoid structure (TLS) formation and anti-tumour immunity such as cDC2, naïve T cells, and cytotoxic NK cells were depleted in S-TDLNs, ALNs, and tumours in metastatic groups, accompanied by an increased infiltration of immunosuppressive macrophages, suggesting impaired immune function (Figure 2J). Notably, while naïve B cells were reduced in metastatic tumours, they were preserved in metastatic LNs, indicating partially retained immune competence in these sites (Figure 2J). We also quantified the expression of lymphoid-structure-associated gene programmes across tissues ^31^. These programmes progressively declined with advancing metastasis, but changes were relatively modest in S-TDLNs (Figure 2K). Altogether, these findings highlight the cellular reprogramming and immune suppression that accompany LN metastasis in TNBC.

### Single-cell Transformer model identifies key immune features during LN metastatic transition

In addition to examining changes in immune cell proportions, we also investigated global gene expression alterations to identify key immune features across different patient tissues. To better capture complex gene–gene interactions underlying immune reprogramming, we developed a Transformer-based machine learning model that encodes both gene identity and expression magnitude to predict cell states with high accuracy. This method begins by preprocessing the scRNA-seq data to remove low-quality cells and genes, followed by normalisation of gene expression and selection of highly variable genes. The gene expression data is then encoded into two distinct types of embeddings: one representing the gene itself, and the other reflecting its expression magnitude. These embeddings are combined into gene tokens, which are processed by a Transformer-based model. The Transformer model facilitates communication between tokens, updating a special classification token (CLS) that represents the entire cell. Finally, this updated CLS token is passed through a multilayer perceptron classifier, which predicts the cell’s status based on gene interactions, leveraging advanced machine learning to deliver a precise prediction of cell characteristics (Figure 3A).

**Figure 3.**
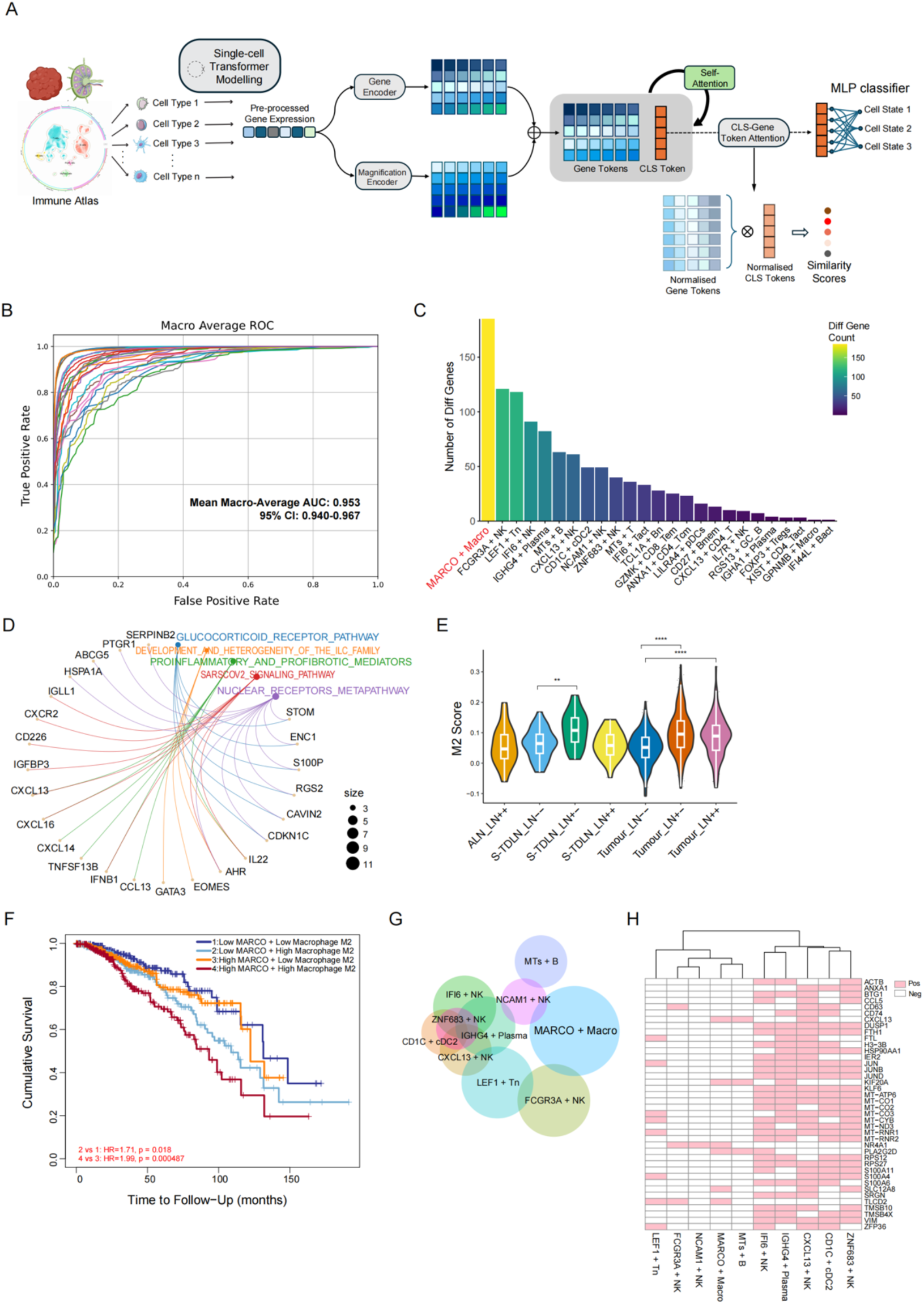
Single-cell Transformer model identifies key immune features driving lymph node metastasis. A, Schematic illustration of the single-cell Transformer-based analytical pipeline. B, Model performance summary across immune cell populations from tumours, mLNs, and uLNs. C, Bar plot summarising the number of significantly differential feature genes identified across immune subsets from different tissues. D, Enrichment analysis of the top-ranked feature genes in MARCO+ macrophages identified through the Transformer model. E, Violin plot comparing immunosuppressive (M2) macrophage scores across lymph node and tumour samples. Boxplots represent median and interquartile range; statistical significance calculated by Wilcoxon test (**P < 0.01, ****P < 0.0001). F, Kaplan–Meier survival curves stratified by MARCO and M2 macrophage gene signatures. Hazard ratios (HR) and log-rank p-values indicated. G, Venn plot illustrating overlapping feature genes identified by the Transformer model across various immune populations. H, Heatmap highlighting selected genes consistently identified across three or more immune subsets.

Using the immune cell subsets from the immune atlas of LN metastasis described in Figure 2, we built separate models for each immune cell population to identify distinguishing feature genes across the tumour, mLN, and uLN groups. The mean area under the curve (AUC) for all immune cell subsets was 0.953, with a 95% confidence interval ranging from 0.940 to 0.967, indicating that key features distinguishing these groups could be identified across many cell types (Figure 3B). We summarised the number of genes considered important features during the self-attention classification process, with MARCO+ macrophages exhibiting the most variable features among all cell types (Figure 3C). Gene enrichment analysis of the top-ranked features in MARCO+ macrophages revealed significant involvement of glucocorticoid signalling, nuclear receptor pathways, and inflammatory processes (Figure 3D), consistent with our previous findings implicating steroid metabolism in TNBC progression ^24^. The M2 macrophage score, representing an immunosuppressive phenotype, was elevated in tumours and LNs of patients with metastatic involvement, suggesting a tumour-promoting reprogramming of the myeloid compartment during LN metastasis (Figure 3E). Notably, we discovered that MARCO expression, along with M2 macrophages, was associated with breast cancer patient survival, underscoring the critical role of MARCO+ macrophages in treatment response and prognosis (Figure 3F).

In addition to MARCO+ macrophages, our Transformer model identified several feature genes associated with both tumours and LNs in FCGR3A+ NK cells and LEF1+ naïve T (Tn) cells, and a substantial overlap of feature genes was found across various NK cell subsets, IGHG4+ plasma cells, and cDC2 (Figure 3G). Heatmap analysis revealed that any feature genes overlapping in more than three immune subsets were prominent, with many mitochondrial-related genes appearing in several cell types, suggesting potential metabolic reprogramming across tissues and during metastasis (Figure 3H). In summary, these results showed that key immune features and transcriptional reprogramming events identified by our Transformer model were associated with LN metastasis in TNBC.

### Disruption of immune cell co-localisation and lymphoid architecture in metastatic lymph nodes

To further explore spatial immune programmes in uninvolved and metastatic S-TDLNs and ALNs, we first performed Visium spatial transcriptomics on three intact LNs representing non-metastatic, intermediate-metastatic, and fully metastatic stages, as confirmed by pathology reports (Figure S3A). Cell types were annotated based on top marker expression (Figure S3B), revealing major immune and non-immune populations (Figure 4A). Within these LNs, immune hubs composed of myeloid, T, and B cells were clearly identified, often were close to epithelial compartments. Cellular composition varied significantly across LN groups, with reduced frequencies of DCs and B cells and increased epithelial and fibroblast populations in metastatic LNs (Figure 4B). We next assessed the capacity for lymphoid structure formation and found a progressive decline in lymphoid formation scores with advancing metastasis, although this reduction was relatively modest in intermediate LNs (Figure 4C). Spatial mapping showed that these scores were highest within immune-rich hubs and lowest in tumour-contacting regions (Figure 4D). Meanwhile, the suppressive M2 macrophage score was significantly elevated in both intermediate and fully metastatic LNs compared to uLNs (Figure 4E), with M2 macrophages localised in both immune hubs and tumour-adjacent areas (Figure 4F).

**Figure 4.**
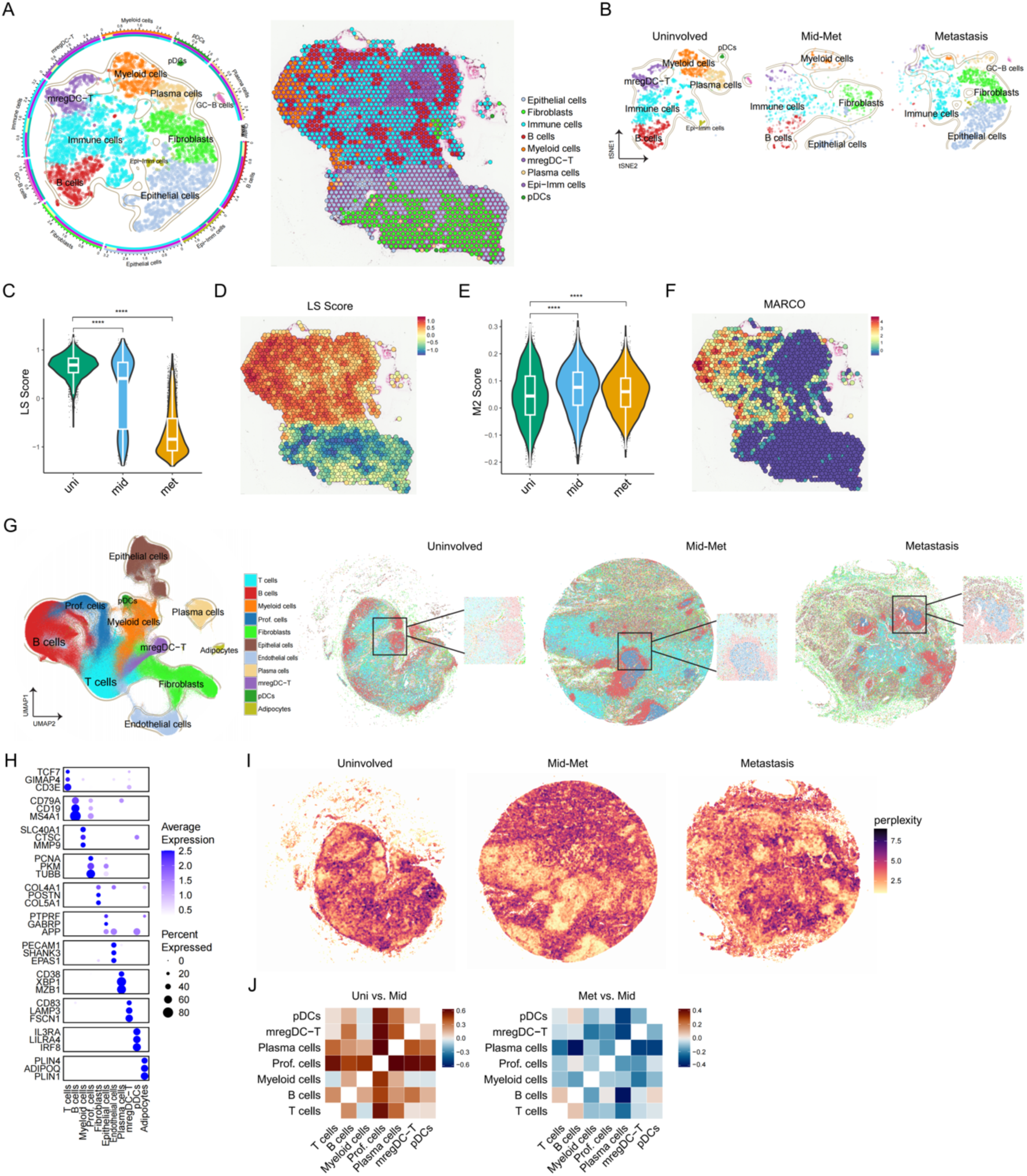
Spatial transcriptomic analysis reveals progressive disruption of immune cell co-localisation and lymphoid architecture during lymph node metastasis. A, Spatial transcriptomics (Visium) profiling of TDLNs across metastatic stages (uninvolved, intermediate-metastatic, fully metastatic). Left, tSNE plot showing major immune and nonimmune cell populations identified based on top marker gene expression. Right, spatial map of intermediate-metastatic TDLN demonstrating annotated cell populations across an intact lymph node sample. B, tSNE visualisation of cellular composition differences across LN groups (uninvolved, intermediate-metastatic [mid-met], metastatic). C, Quantitative comparison of lymphoid structure (LS) formation scores across uninvolved, intermediate-metastatic, and metastatic LNs. D, Spatial distribution of lymphoid structure scores in intermediate-metastatic LN. E, Quantitative comparison of immunosuppressivqe M2 macrophage scores across uninvolved, intermediate-metastatic, and metastatic LNs. F, Spatial mapping of *MARCO* expression in intermediate-metastatic LN. G, High-resolution spatial transcriptomics (Xenium) profiling of TDLNs across different metastatic stages. Left, UMAP plot showing major immune and non-immune cell populations identified based on top marker gene expression. Right, three representative spatial maps of uninvolved, intermediate-metastatic, fully metastatic TDLNs demonstrating annotated cell populations across intact LN samples. Insets indicate regions of significant spatial interactions. H, Dot plot of key marker genes used for annotation of immune and stromal cell types in Xenium data. Dot size represents percentage expression; colour intensity denotes average gene expression. I, Quantitative visualisation of spatial neighbourhood complexity scores across LNs at different metastatic stages. J, Co-localisation heatmaps illustrating pairwise spatial proximity among immune cell subsets, comparing uninvolved vs. intermediate-metastatic (left) and metastatic vs. intermediate-metastatic (right) lymph nodes.

We additionally profiled 13 additional FFPE LNs using Xenium spatial transcriptomics, generating a high-resolution spatial atlas across TDLNs at different metastatic stages (Table S1, Figure S3C). In total, 1,138,735 cells were profiled and classified into major immune and stromal cell types (Figure 4 G,H). Spatial patterns revealed that while T and B cells were preserved in unmetastatic LNs, their organisation became increasingly disrupted in metastatic stages, with epithelial cells encroaching into T and B cell regions (Figure 4G, S3D). We also identified microenvironments enriched for LAMP3+ DCs (mregDCs) in close proximity to T cells, suggesting active regulatory DC–T cell interactions in LNs (Figure 4H). To systematically assess spatial organisation, we quantified neighbourhood complexity across disease stages. In metastatic LNs, spatial connectivity was reduced in B-cell-rich regions and increased near T cells in contact with epithelial zones (Figure 4I). Co-localisation analysis showed that immune–immune cell proximity scores were markedly reduced in S-TDLN+ALN– LNs and further diminished in S-TDLN+ALN+ LNs (Figure 4J). Together, these results reveal stepwise spatial reorganisation and immune disconnection during LN metastasis.

### Multi-modal cell-cell interaction analysis uncovers loss of DC and B–T cell communication during metastasis

To understand how immune cell interactions are altered during LN metastasis, we developed an integrated cell–cell interaction (CCI) analysis framework combining data from scRNA-seq, IMC, and Xenium spatial transcriptomics (Figure 5A). Each modality contributes complementary information: scRNA-seq provides transcript-level ligand–receptor (LR) expression profiles, IMC captures spatial proximity between cell types based on protein-level data, and Xenium enables in situ quantification of RNA expression across intact tissues at single-cell resolution (Figure 5A). We first calculated cell–cell distance metrics from IMC to estimate direct physical contact between immune populations (Figure S4A). In parallel, we applied the CellChat algorithm to scRNA-seq data to predict LR-mediated communication strength between immune cell types based on gene expression patterns (Figure S4B). We then defined an interaction score (IntScore) that integrates spatial proximity from IMC with LR interaction probabilities from CellChat across samples from uLNs, mLNs, and tumours. This allowed us to compare immune communication networks across tissues using harmonised cell-type annotations (Figure 5B).

**Figure 5.**
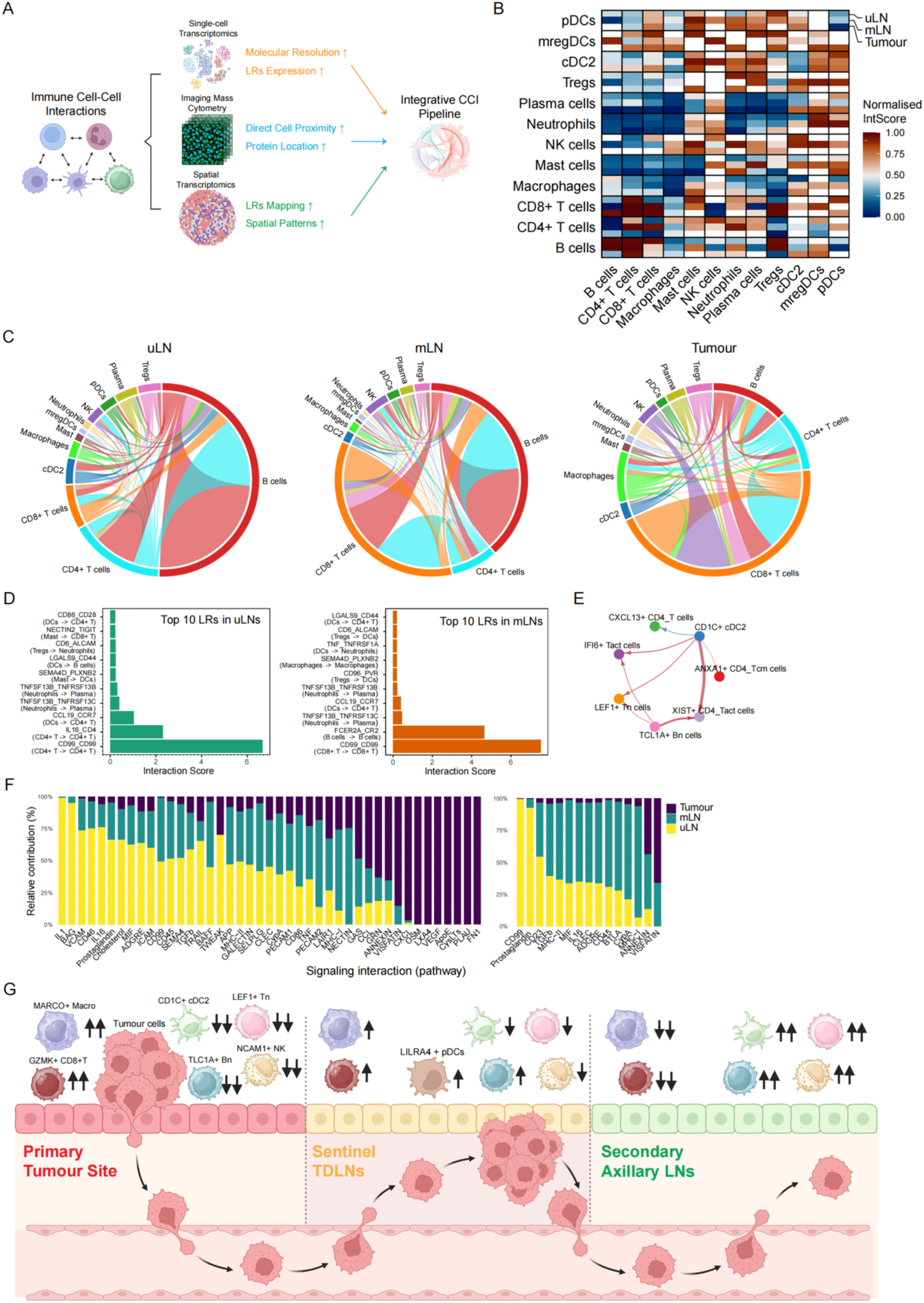
Multi-modal cell–cell interaction (CCI) analysis reveals impaired dendritic cell and B–T cell communication during LN metastasis. A, Overview of the integrated CCI analytical pipeline. The complementary modalities like ligend and receptors (LRs) were integrated to construct comprehensive immune interaction networks across tissues. B, Heatmaps illustrating integrated interaction scores (IntScore) combining IMC-based cell proximity with CellChat-derived LR interaction probabilities for major immune subsets across uLNs, mLNs, and tumours. Lower interaction scores indicate reduced cell–cell interaction possibilities. C, Circular network plots depicting significant LR interactions among immune cell subsets in uLNs, mLNs, and tumour tissues. D, Bar plots summarising the top 10 LRs identifying in uLNs and mLNs. E, Circular network plot visualising cell–cell interaction strengths specifically among DCs, B cells, and various T cell subsets. F, Relative bar plots detailing specific alterations in LR pathways within CD1C+ cDC2 and TCL1A+ naïve B cells in uLNs, mLNs, and tumour tissues. G, The graphical illustration shows how immune population alterations during human sequential LN metastasis.

Visualisation of significant LR pairs among all cell types revealed a reduction in signalling activities involving cDC2, B cells, and CD4+ T cells in tumours compared to uLNs, while macrophages and CD8+ T cells mediated interactions were increased (Figure 5C). Next, we integrated spatially-resolved LR interactions identified through SpatialDM analysis of Xenium data (Figure S4C) with interaction probabilities predicted from scRNA-seq. Investigation of the most significantly altered LR pairs in uLNs and mLNs revealed impaired co-stimulatory interactions and immune-cell recruitment, particularly between DCs and T cells (Figure 5D). To further characterise alterations in immune communication, we analysed the interaction strengths and LR pairs specifically among DCs, B cells, and T cell subsets. This network analysis highlighted substantially weakened interactions in tumour samples, particularly affecting immune subsets involved in antigen presentation and lymphoid structure formation (Figure 5E, S4D).

Given the observed loss of CD1C+ cDC2 and TCL1A+ naïve B cells in metastatic tissues, we further analysed changes in LR pathways specific to these cell types. In cDC2, we found increased activity in IL-1, MHC-II, and CD86 signalling, along with reduced expression of VEGF and ApoE pathways. In naïve B cells, CD23 and MHC-II pathways were elevated, while VISFATIN was decreased (Figure 5F). Together, this multi-modal CCI analysis highlights a consistent disruption of immune communication during LN metastasis, particularly affecting dendritic cell and B cell signalling with T cells. These changes may contribute to the impaired immune surveillance observed in advanced TNBC.

### CD1c⁺ cDC2 cells in lymph nodes predict clinical benefit in TNBC patients receiving neoadjuvant immunotherapy

To evaluate the clinical relevance of our previously identified immune alterations during LN metastasis, particularly the depletion of specific T and B cell clusters and cDC2, we recruited an independent cohort of 36 TNBC patients receiving neoadjuvant chemotherapy plus anti-PD-1 therapy (Table S1, Figure S5A). After 6-8 cycles of treatment, patients underwent surgery and were pathologically classified into pathologic complete response (pCR, n = 18) and non-pCR (n = 18) groups based on residual disease status in the breast and lymph nodes. Multi-colour immunofluorescence (mIF) was performed on surgically resected lymph nodes to assess immune population frequency and its association with treatment response and survival (Figure 6A).

**Figure 6.**
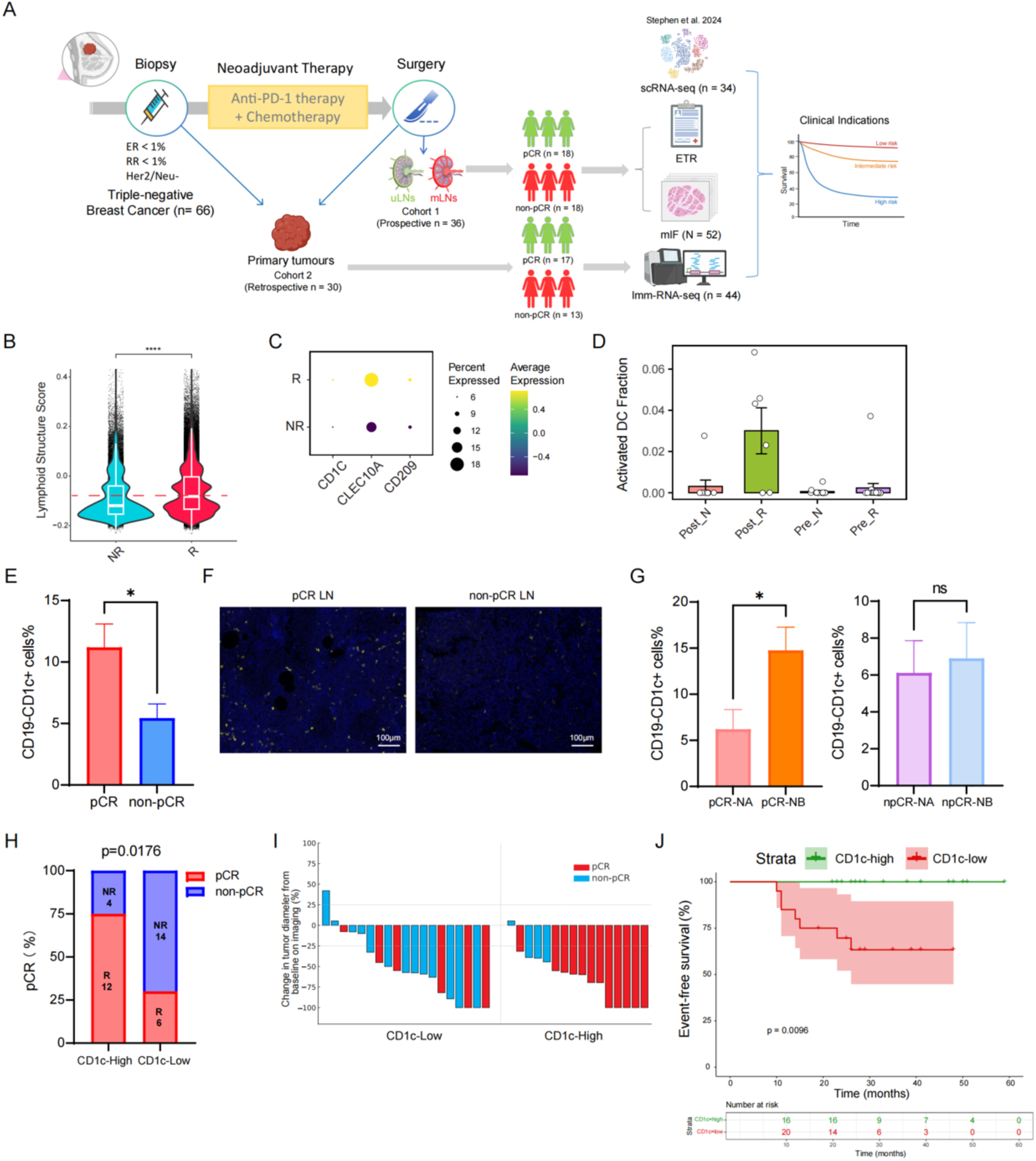
CD1c⁺ cDC2 cells in lymph nodes predict clinical benefit in TNBC patients receiving neoadjuvant immunotherapy. A, Schematic overview of patient recruitment, treatment, and sample processing. A total of 66 TNBC patients receiving neoadjuvant chemo-immunotherapy were enrolled and classified into pCR and non-pCR groups following surgical resection and pathological evaluation. B, Violin plot showing lymphoid structure scores in tumour-infiltrating immune cells from responders (R) compared to non-responders (NR), based on analysis of public scRNA-seq data from 34 TNBC patients. C, Dot plot visualising gene expression of cDC2 markers (CD1C, CLEC10A, CD209) in myeloid cells from responders, based on analysis of public scRNA-seq data from 34 TNBC patients. D, Barplot showing activated DC fraction in responders versus non-responders before and after the treatment, based on RNA-seq data from 44 TNBC tumour samples. E–F, Multi-colour immunofluorescence analysis showing the proportion of CD1c⁺ cDC2 cells in lymph nodes from pCR versus non-pCR patients (E), with representative images from two patients illustrated in (F). G, Multi-colour immunofluorescence analysis showing the proportion of CD1c⁺ cDC2 cells in regressed metastatic nodes (Sataloff score NA) and uninvolved lymph nodes (Sataloff score NB) from pCR (left) and non-pCR patients (right). H, Barplot visualising the proportion of pCR in CD1c-high (≥ average) and low (< average) patients. I, Waterfall plot showing percentage change in tumour diameter from baseline on imaging in CD1c-high (≥ average) and low (< average) patients. J, Kaplan–Meier analysis of event-free survival (EFS) stratified by CD1c⁺ cDC2 status. All seven events (2 locoregional recurrences, 5 distant metastases) occurred in the CD1c-low (< average) group during a median follow-up of 28.5 months. Statistical tests used: two-sided Wilcoxon or unpaired t-test; Kaplan–Meier analysis with log-rank test. *P < 0.05, **P < 0.01, ****P < 0.0001.

To investigate whether immune activities affected by LN metastasis were linked with treatment responsiveness, we first analysed a public scRNA-seq dataset of TNBC patients receiving immunotherapy ^32^. Responders (R) exhibited higher lymphoid structure scores and lower M2 macrophage gene signatures including *MARCO* in tumour-infiltrating immune cells compared with non-responders (NR) (Figure 6B, Figure S5B). Notably, the expression of cDC2 markers (*CD1C, CLEC10A, CD209*) was significantly upregulated in myeloid cells from responder patients (Figure 6C), suggesting a potential role of cDC2 in effective immune responses. In addition, cDCs were also found correlated with breast cancer patient survival (Figure S5C). We further analysed our in-house bulk RNA-seq data from 44 pre-treatment tumour biopsies and surgical samples of 30 TNBC patients (NCT04676997) ^33^. Consistently, the fraction of activated dendritic cells was significantly higher in responders than in non-responders (Figure 6D), indicating the association between dendritic cell presence and therapeutic efficacy.

Focusing on lymph nodes, we found that CD1c⁺ cDC2 cells were significantly more abundant in the LNs of pCR patients than non-pCR patients, as assessed by mIF (Figure 6E,F). Within the pCR group, uninvolved LNs (Sataloff score NB) showed higher percentages of CD1c⁺ cDC2 cells compared to LNs where metastatic components had regressed (Sataloff score NA), and this pattern was not observed in the non-pCR group (Figure 6G, S5D,E). These findings were correlated with our earlier observation that cDC2 were more in treatment-naïve patients without LN metastasis, suggesting that the immunological baseline of the lymph node, particularly in initially uninvolved nodes, played a critical role in determining therapeutic efficacy. To assess clinical impact, patients were classified into CD1c-high and CD1c-low groups based on the percentage of CD1c⁺CD19⁻ cells in LNs. A significantly greater proportion of CD1c-high patients achieved pCR (Figure 6H), and imaging analysis revealed a more pronounced reduction in tumour diameter from baseline in these patients (Figure 6I). Importantly, patients in the CD1c-high group exhibited significantly improved event-free survival (EFS) compared to those in the CD1c-low group; Over a median follow-up of 28.5 months, all seven recorded events, including two cases of isolated locoregional recurrence and five cases of distant metastasis, occurred exclusively in the CD1c-low group (Figure 6J).

Together, these findings demonstrate that cDC2 preservation in LNs during neoadjuvant immunotherapy is associated with improved tumour regression and favourable clinical outcomes. This highlights CD1c as a potential predictive biomarker and therapeutic target, supporting the translational relevance of our immune landscape of LN metastasis and dendritic cell-based interventions for TNBC.

## DISCUSSION

In this study, we presented a comprehensive spatial and transcriptional atlas of immune remodelling in paired primary tumours, sentinel TDLNs, and secondary LNs during the progression of TNBC. By integrating scRNA-seq, spatial transcriptomics (Visium and Xenium), IMC, and mIF, we depicted the stepwise disruption of lymphoid architecture and immune cell interactions associated with LN metastasis. Our findings reveal that early immune reprogramming events in metastatic TDLNs, underscoring the critical role of TDLNs in shaping antitumour immunity.

The spatial and transcriptional immune roadmap constructed in this study highlights the progressive loss of organised lymphoid structures in metastatic LNs and tumour microenvironment. Notably, we observed a decline in key immune cell populations such as cDC2 and naïve T cells, which are essential for initiating and sustaining adaptive immune responses ^34–36^. Previous studies have shown that CD11c⁺ DCs were enriched in TLS-high and TIL-rich primary tumours with abundant CD4⁺ and CD8⁺ T cells, and were associated with improved clinical outcomes in treatment-naive TNBC patients ^37^. In addition, compared with healthy individuals, significantly less naïve T cells were found in blood samples of breast cancer patients, which could be transformed to effective phenotype under the right conditions ^38^. In our study, the depletion of these lymphoid-structure-related populations correlated with impaired antigen presentation and diminished T-cell priming in mLNs with advanced stages, suggesting a compromised immune surveillance mechanism in metastatic settings.

The application of our single-cell Transformer model enables the identification of key immune features and cell states associated with LN metastasis. Transformer-based approaches have increasingly been employed to extract biological insights from the complex processes underlying cancer biology and immunology ^39–41^. By integrating advanced machine learning with high-dimensional single-cell data, our model effectively captured intricate gene expression patterns and identified key feature genes across diverse immune cell types, providing insights into the dynamic immune landscape of TNBC LN metastasis. Notably, the model highlights the elevated proportion of MARCO⁺ macrophages in metastatic LNs and tumour tissues from patients with advanced nodal stages, indicating their role in tumour progression and immune evasion. The presence of MARCO⁺ macrophages, known for their immunosuppressive functions, has been associated with tumour growth and metastasis, and serve as a potential therapeutic target ^42–44^. Our findings further characterise their immunosuppressive profile in the context of LN metastasis, suggesting that therapeutic strategies targeting MARCO⁺ macrophages within both metastatic LNs and primary tumours may improve immunotherapy efficacy in TNBC.

Furthermore, our integrative cell-cell interaction (CCI) analysis framework, which combines scRNA-seq, IMC, and Xenium spatial transcriptomics, uncovered the disruption of coordinated immune spatial architecture in TNBC LNs and tumours (Figure 5). We observed a significant loss of dendritic cell and B–T cell communication during LN metastasis, particularly affecting cDC2 and naïve B cells. Cross-talk between DCs and T cells in TDLNs has been shown to be crucial for breaking immune tolerance ^45^, and its disruption has been linked to immune evasion, tumour progression, and poor clinical outcomes in breast cancer ^46^. Similarly, specific T–B cell interactions in primary TNBC tumours have been shown to carry significant prognostic value ^47^. Our findings demonstrated the breakdown of DC/B-T networks within metastatic TDLNs contributed to immunosuppressive microenvironment, facilitating tumour progression. Therapeutic strategies aimed at restoring these interactions—such as the use of DC vaccines ^48^ may represent a viable strategy to bolster antitumor immunity, particularly when combined with anti-MARCO+ macrophage strategies ^42^.

Clinically, our study is highly related to the evolving paradigm of axillary management in breast cancer. Sentinel lymph node biopsy (SLNB) has emerged as a milestone in breast surgery, marking a shift towards minimally invasive approaches that preserve TDLNs ^11^. Recent trials, such as SENOMAC and INSEMA, have demonstrated the safety of omitting completion axillary lymph node dissection in selected patients, highlighting the feasibility of surgical de-escalation ^14,17^. However, Traditional surgical strategies have primarily focused on the mechanical assessment of tumour cell metastasis and balancing oncological safety and quality of life, often overlooking the immunological significance of LNs. Our findings advocate for a better understanding of TDLNs as active immunological hubs, whose preservation may be critical for maintaining systemic antitumor immunity. This perspective is particularly pertinent in TNBC, where the efficacy of immune checkpoint inhibitors remains limited compared with hot tumours like lung cancer ^49^. By recognising the immunological functions of TDLNs, we can better determine optimal windows for immunotherapeutic intervention and improve patient outcomes—for example, by administering immunotherapy prior to surgery ^50^. In support of this, we found that TNBC patients with preserved CD1c⁺ cDC2 in lymph nodes following neoadjuvant immunotherapy exhibited significantly higher rates of pathological complete response and improved event free survival, highlighting the prognostic and therapeutic relevance of immune cell composition in preserved nodes.

Together, our study provided a comprehensive spatio-temporal transcriptional map of immune remodelling across primary tumours, sentinel, and non-sentinel lymph nodes in treatment-naive TNBC. We revealed early depletion of immune populations related to lymphoid architecture, impaired cell–cell communication, and enrichment of immunosuppressive compartments during LN metastasis (Figure 5G). Future work should investigate whether the immune features we identified during LN metastasis possess prognostic value for current immunotherapies, and, more importantly, how combining anti-MARCO+ macrophage strategies with DC vaccines might be integrated into treatment regimens to improve outcomes in this challenging breast cancer subtype.

## METHODS

### Human specimens and ethical approval

This study enrolled 116 female patients diagnosed with early-stage or locally advanced triple-negative breast cancer (TNBC) at the Shandong Cancer Hospital and Institute (Jinan, China). The cohort included 50 patients who underwent upfront surgery and 66 patients who received neoadjuvant therapy, and total of 243 samples were collected.

#### Upfront Surgery Cohort

women with TNBC who planned to undergo upfront surgery were eligible for the study if they were 18-75 years of age and had a clinical tumour stage of T1 or T2 (tumor size, ≤5 cm) and node-negative status according to clinical assessment (cN0) (imaging abnormalities were allowed). Exclusion criteria included ongoing pregnancy or breast-feeding, inflammatory breast cancer, in situ breast cancer, bilateral breast cancer, distant metastases, prior malignancy, neoadjuvant therapy, disease or comorbidity possibly affecting the immune status, and obstacles to obtaining informed consent. A total of 50 female patients were enrolled, with 147 specimens collected and analysed: including 48 primary tumour samples and 99 regional lymph node samples.

#### Neoadjuvant Therapy Cohort

sixty-six patients with TNBC (clinical stage cT0-1cN1-3M0 or cT2-4N0-3M0) received neoadjuvant therapy (30 cases were retrospectively collected from our previous clinical trial, NCT04676997), typically consisting of 6-8 cycles of neoadjuvant immunotherapy (PD-1 inhibitors) combined with chemotherapy, following institutional guidelines. Exclusion criteria included inflammatory breast cancer, Incomplete neoadjuvant therapy, bilateral breast cancer, special types breast cancer, prior malignancy, disease or comorbidity possibly affecting the immune status and ongoing pregnancy or breast-feeding. A total of 44 primary tumour samples and 52 regional lymph node samples were collected from this cohort.

TNBC was defined as human epidermal growth factor receptor 2 (HER2)-negative (immunohistochemistry [IHC] score 0 or 1+, or IHC 2+ with negative fluorescence in situ hybridisation [FISH]) and oestrogen receptor (ER)- and progesterone receptor (PR)-negative (<1% expression). The study protocol was approved by the Ethics Committee of Shandong Cancer Hospital and Institute, affiliated with Shandong First Medical University and Shandong Academy of Medical Sciences (approval no. SDTHEC2023009011). Written informed consent was obtained from all participants, allowing for tissue collection and downstream analyses.

### Sample collection

#### Upfront Surgery Cohort

Procedures included mastectomy (n = 38) or lumpectomy (n = 12), and all underwent SLNB. Sentinel lymph nodes were defined as any lymph nodes with a standard accumulation of tracer substances (blue dye or isotopes) or suspected of being cancerous by intraoperative palpation. Lymph nodes that did not fulfill these criteria but that had been randomly removed during surgery were not classified as sentinel lymph nodes. If the sentinel lymph node contained a metastasis, complete axillary dissection was performed immediately. Axillary lymph node dissection was defined as a dissection of at least anatomical levels I and II including at least ten nodes. Nodes collected via axillary lymph node dissection or unqualified as sentinel lymph node were termed ALNs in this study. The recommendations of the American Joint Committee on Cancer were applied for the assessment of histopathological specimens. Intraoperative assessment involved sectioning all S-TDLNs and one ALN into 2-mm blocks for touch imprint cytology and frozen sectioning. Fresh tumour, S-TDLN, and ALN samples were immediately preserved. Postoperatively, all nodes underwent standard histopathological evaluation. A lymph node was classified as positive if any of cytology, frozen section, or routine pathology identified metastasis. Based on S-TDLN/ALN status, patients were grouped as: S-TDLN−/ALN− (n = 20), S-TDLN+/ALN− (n = 15), or S-TDLN+/ALN+ (n = 15).

#### Neoadjuvant Therapy Cohort

Surgical and pathological assessment procedures mirrored those of the initial surgery cohort. Total pathological complete response (pCR) was defined as the absence of residual invasive carcinoma in the primary breast tumor (ypT0/is) and no metastatic cancer in the ALNs (ypN0). The pathological states of ALNs after NAT were classified as follows N-A: (1) No metastatic cancer cells present, but post-treatment changes observed (indicating prior involvement). (n=18 specimens) (2) N-B: No metastatic cancer cells present, and no post-treatment changes observed. (n=28 specimens) (3) N-C: Metastatic cancer present, along with post-treatment changes. (n=5 specimens) (4) N-D: Metastatic cancer present, without post-treatment changes. (n=1 specimen)

### Histological staining and tissue microarray (TMA) construction

Formalin-fixed, paraffin-embedded (FFPE) blocks were prepared after fixation in neutral-buffered formalin for 24–48 hours, followed by graded ethanol dehydration, xylene clearing, and paraffin embedding at 60°C. Hardened blocks were sectioned into 4-μm slices, floated in a 37°C water bath, mounted onto charged slides, and baked at 65°C for 2 hours. H&E staining followed standard procedures: xylene dewaxing, ethanol rehydration, Harris haematoxylin staining (5–10 min), acid alcohol differentiation, bluing, eosin counterstaining (1–3 min), and dehydration/clearing. Coverslipped slides were evaluated by light microscopy.

IMC, Xenium Spatial Transcriptomics, and multiplex immunofluorescence (mIF) assays were performed using TMAs. For TMA generation, immune-rich regions were selected from H&E slides. Tissue cores with diameters of 3.0 mm (for IMC and Xenium) or 2.0 mm (for mIF) were then extracted from these regions and arrayed into a recipient paraffin block.

### Single-cell RNA sequencing and analysis

Fresh tissues were rinsed in cold PBS, minced into ∼1 mm³ pieces, and incubated in PBS containing DNase I (100 μg/mL) for 5 minutes at room temperature. Mechanical dissociation was achieved via 100-μm strainers and syringe plungers. Cell pellets were lysed with RBC Lysis Buffer, washed with PBS + 0.04% BSA, and filtered. CD45⁺ cells were enriched using fluorescent antibodies and sorted on a BD FACSAria III (70-μm nozzle), yielding >95% purity. Viability was assessed via trypan blue staining on a Countess II FL counter. Samples with >85% viable cells proceeded to library prep; otherwise, dead cells were removed using a Miltenyi Biotec kit.

Libraries were generated using the 10X Chromium Single Cell 3′ v3 platform, targeting 8,000 cells/sample. Reverse transcription and barcoding occurred in GEMs. Libraries were amplified, fragmented, adaptor-ligated, and enriched. QC was performed via Agilent Bioanalyzer. Sequencing was done on Illumina NovaSeq 6000 (150 bp PE), aiming for ≥20,000 reads/cell. Raw data were processed using Cell Ranger (v7.0.0) and aligned to GRCh38. Filtering excluded cells with < 200 or > 7,500 genes, > 50,000 UMIs, or > 10% mitochondrial reads. Seurat (v5.0.3) ^51^ was used for log-normalisation, PCA, clustering, and the Uniform Manifold Approximation and Projection (UMAP) or t-Distributed Stochastic Neighbor Embedding (t-SNE). A total of 246,703 cells passed QC and were annotated based on their marker gene expression.

To infer intercellular communication networks from scRNA-seq data across different tissue compartments, we employed the CellChat (v2.1.2) R package ^52^. For different tissue datasets, raw UMI count matrices and associated cell type metadata were used to construct individual CellChat objects, applying the human ligand–receptor interaction database (CellChatDB.human). Following standard workflows, overexpressed genes and interactions, computed communication probabilities and pathway-level signaling strength, and evaluated network centrality metrics were identified. The outputs were visualised included ranked pathway strength and source–target flow analyses.

### Single-cell Transformer model

#### (1). Preprocessing of raw counts single-cell RNA-seq data

The preprocessings are performed using the scanpy package in Python. Given a batch of raw count single cell RNA (scRNA) 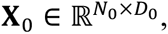 with N being the number of cells, *D*_0_ being the original number of available genes. **X**_0_ goes through several procedures to reach the final product, **X** ∈ ℝ*^N^*^×*D*^ where *N* and *D* are the numbers of filtered cells and genes, respectively. The procedures include:

1. Quality Control: This step is to exclude unqualified cells and genes. Unqualified cells refer to those with unusual total number of detected genes, or those with mitochondrial gene expression being higher than a threshold. Meanwhile, the genes expressed in very minor cells (e.g., less than 5 cells) are regarded as unqualified genes.
2. Normalisation: an scRNA sequence is normalised and scaled so that the total expression is summed up to a fixed value (e.g., 1e4). Then it is transformed by the log1p operation.
3. Highly variable genes (HVGs) selection: A fixed number of genes with the highest variations are selected, while the other ones are abandoned as they are seen as being less informative when with lower variations. In the setting, we remain 2000 HVGs.
4. Batch effect correction: Each patient’s scRNA is seen as a batch for the batch effect correction.

#### (2). Method overview

In this section, our model developed to predict cell status is elaborated. Given a pre-processed scRNA sequence with length *D*, each gene in it will be encoded into two types of embeddings by two different encoders, respectively, namely, the gene embedding 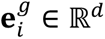 and the expression magnitude embedding 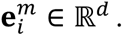 The gene token is then formulated by summing the gene embedding and expression magnitude embedding of an individual gene. A CLS (class) token is introduced, which has the same dimension as the gene token and is learnable during the training phase. All the tokens (gene tokens and CLS token) are forwarded in a series of Transformer-based self-attention modules, followed by a CLS similarity module. Finally, the updated CLS token is fed into an MLP (multilayer perceptron) classifier for the final prediction of cell status. The cosine similarity values derived from the CLS similarity module contribute directly to the downstream prediction, thus they reflect the relevance of genes to the cell status.

#### (3). Genes to gene tokens

We adopt the solution from ^53^ to encode the gene expression with a non-zero scalar value in a scRNA. In that, the gene encoder will transform a particular gene (regardless of its expression level) into a fixed value vector 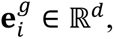 termed the gene embedding. Similarly, another encoder transforms the value of the gene into the expression magnitude embedding 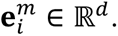 Note that these two encoders are pretrained from ^53^. The gene token **e***_i_* ∈ ℝ*^d^* is generated by,

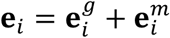

#### (4). Transformer-based self-attention

The self-attention based on Transformer allows mutual-information exchange among tokens.

The learnable CLS token **e***^cls^* ∈ ℝ*^d^* is introduced, which serves as the cell representation after being updated. Let **Z** ∈ ℝ^(*M*+1)×^*^d^* stack **e***_i_* and **e***^cls^*, with *M* being the number of genes of a scRNA,

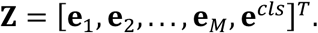

For self-attention operation, **Z** is first projected into queries (**Q**), keys (**K**), and values (**V**),

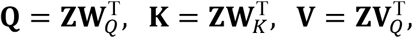

where **W***_Q_*, **K***_Q_*, **V***_Q_* ∈ ℝ*^d’^* ^×*d*^ are learnable weight matrices, and **Q**, **K**, **V** ∈ ℝ*^M^*^×*d*^. The updated token is obtained by,

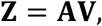

where **A** ∈ ℝ*^M^*^×*M*^ is the correlation matrix and is obtained from the following operation,

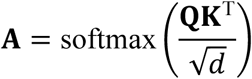

This process repeats for several rounds so that the tokens can exchange information among themselves sufficiently.

#### (5). CLS similarity module

CLS similarity module serves to aggregate information from all gene tokens to the CLS token, with the latter being treated as the cell representation for the cell-level downstream analysis. The aggregation operation is based on the cosine similarity between the CLS tokens with each of the gene tokens, that is,

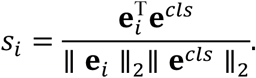

The updated CLS token (cell representation) is obtained by,

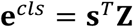

with **s** = [*s*_1_, *s*_2_, … *s_M_*, 1]^T^ ∈ ℝ*^M^*^+1^. A classifier is then imposed on the final CLS token for the final prediction of cell status. Clearly, this operation refines the CLS tokens by aggregating information from the gene tokens based on correlations. Using cosine similarity as the correlation metric, the contribution of each gene token is normalised to be between (-1,1), allowing a standardised comparison.

The final prediction is implemented by an MLP-based classifier,

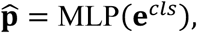

where **p̂** ∈ ℝ^3^ are the estimated probabilities of cell status.

### Visium spatial transcriptomics

Fresh-frozen sections (10 μm) were mounted on Visium slides, stained with H&E, and imaged. Spatial barcoding, cDNA synthesis, and library construction followed the Visium protocol. Libraries were QCed and sequenced (Illumina NovaSeq 6000, 2×150 bp, ≥50,000 reads/spot).

Space Ranger (v2.0.0) and STAR (v2.7.10a) were used for alignment and UMI counting. Spots with <500 UMIs or >25% mitochondrial reads were excluded. Seurat and SpatialLIBD (R) were used for normalisation, clustering, and differential expression (adjusted P ≤ 0.01, |log₂FC| ≥ 0.5). Spatial patterning was analysed using Moran’s I and permutation tests.

### Xenium in situ spatial transcriptomics

FFPE sections (5 μm) were mounted on Xenium slides, deparaffinised, rehydrated, and permeabilised. Probes from the Xenium Prime 5K panel were hybridised overnight at 50°C. Ligations were performed at 37°C. RCA was used for signal amplification. Gene decoding involved iterative hybridisation with labelled probes on the Xenium Analyser, with DAPI-based cell segmentation. Expression matrices were generated. Slides were washed, then stained using Gill II haematoxylin and Eosin Y, dehydrated, and coverslipped. Images were acquired at 40× on a KF-PRO-005 scanner.

Scanpy (v1.9.3), Seurat (v5.0.3), and hoodscanR (v1.0.0) ^54^ used for QC and normalisation. Cells with transcript counts outside the 2nd–98th percentile were excluded. Batch correction was performed using BBKNN. Clustering and visualisation used PCA plus UMAP/t-SNE with Louvain.

Spatial transcriptomics data were analysed to identify cell-cell interactions via ligand-receptor spatial co-expression using SpatialdDM (v0.3.1)^55^. All 1,138,735 cells from 13 samples were processed using common parameters and compared among each other using the differential analysis module. For each sample, a spatial weight matrix was computed using ‘sdm.weight_matrix’ (l=20), followed by extraction of human ligand-receptor pairs with a minimum expression filter of 3 cells using ‘sdm.extract_lr’. Global spatial autocorrelation was assessed via 1,000 ‘sdm.spatialdm_global’ z-score method; and statistically significant ligand-receptor pairs were identified using FDR correction at a 0.1 threshold.

### Imaging mass cytometry

FFPE sections (5 μm) were retrieved in EDTA buffer (pH 9.2), blocked with BSA, and stained with 37 metal-tagged antibodies overnight. Nuclei were labelled with an iridium intercalator. Laser ablation was performed using the Hyperion Imaging System (1 μm² resolution). CellProfiler (v4.1.2) was used for segmentation, and histoCAT or custom scripts for quantification. Batch effects were corrected via Harmony. PhenoGraph or FlowSOM identified cell clusters. Cytomapper and imcRtools (R) were used for spatial analyses. Significance was assessed via Wilcoxon tests and spatial metrics (P < 0.05).

### Integrated cell–cell interaction (CCI) analysis framework

To cross-validate inferred ligand–receptor (L–R) signaling events across immune compartments, we integrated CellChat-derived predictions with two spatial platforms. First, CellChat interaction probabilities based on scRNA-seq data were combined with cell–cell contact frequency data from imaging mass cytometry (IMC). For each sender–receiver cell-type pair, the number of spatially observed interactions were calculated and derived a composite interaction score (IntScore) by multiplying z-scaled CellChat communication probabilities and IMC proximity frequencies. This enabled ranking of spatially supported L–R communications across tissues.

Second, we incorporated L–R interaction scores derived from spatial transcriptomics datasets, where interaction significance was defined by spatial co-localisation metrics (mean z-scores and FDR) for each L–R pair across patient samples. Average spatial z-scores per L–R pair were summarised and merged with CellChat predictions to calculate a joint spatial–transcriptomic score (joint_score = scaled(CellChat) × scaled(spatial z)). For each tissue, the top 10 L–R pairs were identified by their joint scores, highlighting spatially enriched and transcriptionally supported immune communication axes.

### Multiplex Immunofluorescence

FFPE sections (3 μm) were stained using the AlphaTSA multiplex kit. Antigen retrieval (pH 9.0, 95°C, 20 min), blocking, sequential antibody incubation, HRP-conjugated secondary binding, and XTSA dye amplification were performed. DAPI was used for counterstaining. Images were acquired on a PathScan slide scanner. InForm software was used for spectral unmixing, and HALO (Indica Labs) for quantitative cell analysis. Cell densities were computed across tissue compartments, including single- and double-positive immune populations.

## Acknowledgements

The Figure was partly generated using BioRender and Servier Medical Art, licensed under a Creative Commons Attribution 3.0 unported license AI.

## Author contribution

Q.Z. and P.Q. conceptualised and designed the study. Q.Z. analysed the data, and wrote the manuscript. Y.L. and Z.S. collected clinical samples and performed the experiments. H.Z., L.C.S performed part of bioinformatics analyses. R.Z., Y.G., Z.Z., X.S., Y.Q., Y.L., X.W., C.W., B.C., X.N., Y.L., and M.Z. provided assist for experiments. P.Q., B.M., and Y.W. supervised the study.

## Data and materials availability

All data and code used in the analysis will be made available in a public database before the paper is formally accepted.

